# Bioaugmented sand filter columns provide stable removal of pesticide residue from membrane retentate

**DOI:** 10.1101/2020.06.04.135582

**Authors:** Lea Ellegaard-Jensen, Morten Dencker Schostag, Mahdi Nikbakht Fini, Nora Badawi, Alex Gobbi, Jens Aamand, Lars Hestbjerg Hansen

**Affiliations:** Department of Environmental Science, Aarhus University, Roskilde, Denmark; Department of Geochemistry, Geological Survey of Denmark and Greenland (GEUS), Copenhagen K, Denmark; Section of Chemical Engineering, Department of Chemistry and Bioscience, Aalborg University, Esbjerg, Denmark; Department of Plant and Environmental Sciences, University of Copenhagen, Frederiksberg, Denmark

**Keywords:** Drinking water, membrane separation, groundwater contamination, bioremediation, *Aminobacter* sp

## Abstract

Drinking water resources, such as groundwater, are threatened by pollution. The pesticide metabolite 2,6-dichlorobenzamide (BAM) is one of the compounds frequently found in groundwater. Studies have attempted to add specific BAM-degrading bacteria to sand-filters at drinking water treatment facilities. This biotechnology has shown great potential in removing BAM from contaminated water. However, the degradation potential was formerly lost after approximately 2-3 weeks due to a decrease of the degrader population over time.

The aim of the present study was to overcome the constraints leading to loss of degraders from inoculated filters. Our approach was threefold: 1) Development of a novel inoculation strategy, 2) lowering the flowrate to reduce washout of cells, and 3) increasing the concentration of nutrients hereunder the pollutant in a smaller inlet water stream. The two latter were achieved via modifications of the inlet water by applying membrane treatment which, besides producing an ultra-pure water fraction, produced a residual water stream with nutrients including BAM concentrated in an approximately 10-fold reduced volume. This was done to alleviate starvation of degrader bacteria in the otherwise oligotrophic sand-filters and to enable a decreased flowrate.

By this approach, we achieved 100% BAM removal over a period of 40 days in sand-filter columns inoculated with the BAM-degrader *Aminobacter* sp. MSH1. Molecular targeting of the degrader strain showed that the population of degrader bacteria persisted at high numbers throughout the sand-filter columns and over the entire timespan of the experiment. 16S rRNA gene amplicon sequencing confirmed that MSH1 dominated the bacterial communities.

**IMPORTANCE:** Many countries rely partly or solely on groundwater as the source of drinking water. Here groundwater contamination by pesticide residues poses a serious threat to the production of high quality drinking water. Since scarcity of clean groundwater may occur in progressively larger areas both locally and globally, the need for efficient purification technologies is growing. This study shows that a novel system combining membrane treatment and bioaugmented sand-filters can efficiently remove pesticide residues in laboratory columns when applying specific inoculation and flow conditions. Once upscaled, this system can be used directly for pump-and-treat of contaminated groundwater wells or at drinking water treatment plants.

## INTRODUCTION

Groundwater contamination by pesticide residues poses a grave threat to production of high quality drinking water. Drinking water treatment plants (DWTPs) are, when the concentrations exceed the EU threshold limit of 0.1 µg l^-1^, often forced to the costly procedure of closing the affected drinking water abstraction wells and establishing new wells elsewhere. This issue is a problem, especially in countries that rely partly or solely on groundwater as the source of drinking water. Since scarcity of clean groundwater may occur in progressively larger areas, the need for efficient purification technologies is growing.

The pesticide metabolite 2,6-dichlorobenzamide (BAM) is one of the most frequently found groundwater contaminants and in many cases in concentrations exceeding the EU threshold limit for drinking water (e.g. Johnsen 2015, Porazzi et al. 2005, Pukkila and Kontro 2014, Vandermaesen et al. 2016). BAM-pollution can be removed from abstracted groundwater by sorption to granular activated carbon or by membrane filtration. However, these technologies have their limitations and drawbacks (Chern and Chien 2002, Sabio et al. 2004, Sombekke et al. 1997), especially with regards to the energy demand e.g. for regeneration of activated carbon. Although reverse osmosis membranes have been proven to be efficient in removal of BAM from groundwater (Nikbakht Fini et al. 2019), the treatment results in a residual water stream (retentate) with concentrated BAM and other substances as e.g., nutrient salts and organic carbon. This retentate water may encompass around 10% of the total water stream, which is disposed as an undesired waste product.

Alternatively, bioaugmentation of sand filters at DWTPs were suggested as a technology to purify BAM polluted drinking water. Research into this area has shown that the BAM-degrading *Aminobacter* sp. MSH1 can successfully degraded BAM in bioaugmented sand filters (Albers et al. 2015b, Ellegaard-Jensen et al. 2016, Horemans et al. 2016). However, a decline in the number of added BAM-degrading bacteria, accompanied by a decline in overall BAM degradation, was observed within 2-4 weeks of inoculation (Albers et al. 2015b, Horemans et al. 2016). In these studies, several factors were hypothesised to contribute to the loss of inoculated cells from the sand filters including starvation due to too low levels of BAM and assimilative organic carbon (AOC), mass-transfer limitations, antagonism by indigenous microorganisms in the filters (e.g. predation) or simply continuous washout from the filter (Albers et al. 2015b, Horemans et al. 2016).

We combined membrane separation and bioaugmented sand filter treatment to achieve a superior purification system compared to either of the two treatments alone. The advantages being that the residual water stream from the membrane treatment, composed of high concentrations of minerals, AOC and BAM retained by the membrane, is not discarded as waste but instead used as feed to a bioaugmented sand filter. The hypothesis being that the elements in the retentate water will alleviate starvation of the bioaugmented bacteria in the sand filters prolonging their longevity. It was recently shown, in batch experiments, that membrane retentate water stimulated BAM mineralisation by *Aminobacter* sp. MSH1 compared to untreated water (Hylling et al. 2019).

This study aims to investigate if the crucial survival and performance of the BAM-degrader *Aminobacter* sp. MSH1 can be prolonged beyond what has previously been reported, by adding it into laboratory sand filters with membrane residual water as feed. Further, this study developed and used a novel strategy for inoculation of the degrader bacteria in sand filters, which together with the retentate water feed is hypothesized to stimulate better adherence and biofilm formation of the inoculated bacteria in the sand filters – ultimately ensuring a continuous persistence of the introduced bacterial cells in the sand filters.

## MATERIAL AND METHODS

### Chemicals, media and filter material

Analytical-grade 2,6-dichlorobenzamide (BAM; CAS no. 2008-58-4; 98.5 % purity) was acquired from Dr. Ehrenstorfer (Augsburg, Germany) Analytical-grade 2,6-dichlorobenzamide-3,4,5-d3 (BAM-D3; CAS no. 1219804-28-0; 98.2 % purity), used as internal standard for LC-MS/MS method, was acquired from Chiron (Trondheim, Norway). Mineral salt minimal medium (MS) (Sørensen and Aamand 2003) supplemented with 0.24 g l^-1^ (NH_4_)_2_SO_4_, 0.05 g l^-1^ KNO_3_, and 2 g l^-1^ glycerol (MSNC) was used for pre-growth of *Aminobacter* sp. MSH1 (Albers et al. 2014) and as initial feed to the columns. For protozoan enumeration, Neff’s amoeba saline medium with 0.3 g l^-1^ Tryptic Soy Broth (TSB, Oxoid, Hampshire, England) was prepared according to Rønn et al. (1995). The media were sterilized prior to usage. The column material Nevtraco^®^ was supplied by EuroWater (Silhorko, Denmark) and sieved (< 2 mm), washed and sterilised (autoclaved) before use.

### Influent water treatment

The influent water used in the column experiments was collected from treated water reservoir of a DWTP in south-western part of Denmark, Varde (Lerpøtvej Waterworks, DIN Forsyning, south-west Jutland). Comprehensive water analyses were conducted by an accredited company (Eurofins, Miljø A/S) and the results are presented in Table 1. The collected water was subsequently concentrated using a low pressure reverse osmosis (LPRO) XLE membrane in a lab-scale membrane separation system, M20 TestUnit (Alfa Laval, Denmark) as depicted in an earlier study (Hylling et al. 2019).

**Table 1.**
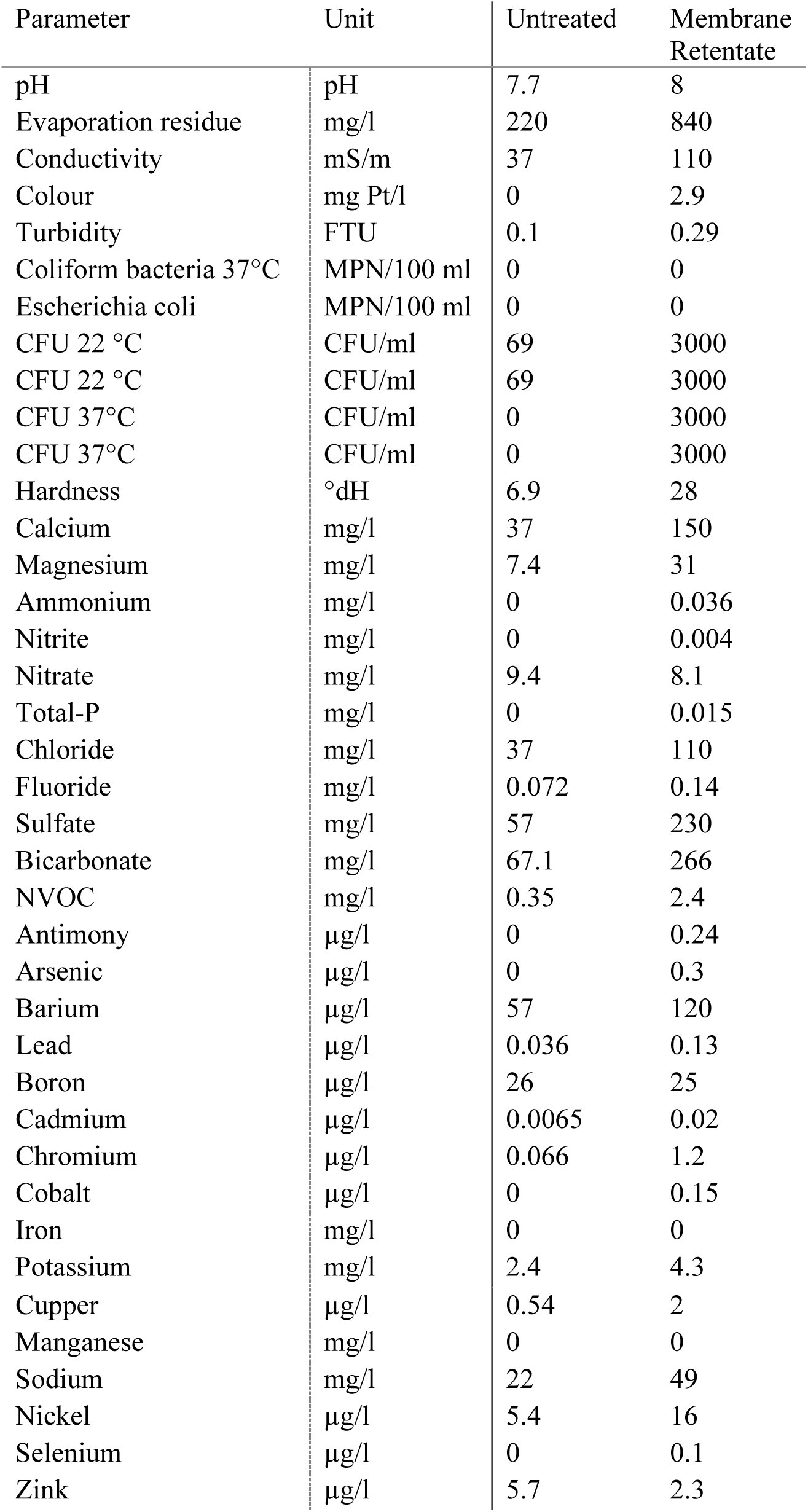
Characterization of the untreated DWTP water and membrane retentate water.

The following protocol was used to produce membrane retentate water samples to be introduced as influent to column sand filters. 25 l of collected water was transferred to the feed tank of the membrane test unit. The feed water was allowed to be recirculated for an hour in a cross-flow mode through a membrane housing hosting the spiral wound XLE membrane with an active area of 1.2 m^2^ in order to reach to the steady state in terms of membrane saturation and both permeate and retentate streams were sent back to the feed tank. The required 5 bar pressure was provided by a Rannie Lab 16.50 reciprocating pump while flow rate was fixed at 8 l min^-1^. The temperature was maintained at 15 °C using a built-in heat exchanger connected to a recirculating chiller. After preconditioning of the membrane, the permeate was collected in a separate container while the retentate was led back to feed reservoir. Membrane filtration was pursued until 90% of the volume of the initial feed water (22.5 l) was obtained as the permeate resulting in a remaining 10× concentrated retentate in the feed tank. The retentate was then collected and stored overnight in sterile bottles at 4 °C and shipped the next day in a cool box to the laboratory for column experiments. Water collected at the same time, but not undergoing membrane treatment was likewise kept cooled and shipped together with the membrane retentate water. The entire experimental protocol was repeated weekly for 6 weeks to produce continuous adequate inlet water for the column experiments.

### Bacterial strain and cultivation

BAM-degrading strain *Aminobacter* sp. MSH1 originally isolated and described by Sørensen et al. (2007) was applied in the present study. MSH1 showed great potential for BAM degradation of membrane retentate water in batch studies (Hylling et al. 2019) and it was recently characterized by full genome sequencing (Nielsen et al. 2018).

The strain was pre-grown in MSNC medium for five days at 20 °C on an orbital shaker (100 rpm). At cell harvest, growth was measured by OD_600nm_ and cells were centrifuged (8000×g for 10 min), washed and re-suspended in 40 ml MSNC containing 10^9^ cells ml^-1^ further use in column experiments.

### Bioaugmented sand filter columns

Laboratory columns experiments were conducted in glass columns (h20 x Ø1.6 cm; GE Healthcare, Thermo Fisher Scientific Inc., Roskilde, Denmark) each wet packed with 50g Nevtraco material. The glass columns had been sterilised with 0.01 M NaOH and rinsed with sterile MilliQ water prior to use. The novel strategy for inoculation was developed to improve adherence of the MSH1 strain to the sand columns, including the following steps: 1) six of the columns were inoculated with 5 ×10^9^ MSH1 cells each by pumping the pre-grown culture of *Aminobacter* sp. MSH1 into the columns, 2) the culture was left in the columns without flow for two hours in order for the cells to adhere to the filter material (Albers et al. 2014), and 3) MSNC was applied as initial feed to the columns for the 48 hours following inoculation. This was done to circumvent the degrader depletions seen in previous experiments (Albers et al. 2015b, Horemans et al. 2016). Two additional columns served as non-inoculated controls. The columns were hereafter supplied with a continuous upward flow of 4.8 ml h^-1^. After the initial period the feed was changed to water collected from Varde DWTP; supplying four columns (three inoculated and one control) with membrane retentate water spiked with 3.0 µg l^-1^ BAM and the four other columns (three inoculated and one control) with untreated water spiked with 0.3 µg l^-1^ BAM. The experiment was carried out at 10 °C for 40 days.

Sampling of outlet water from the individual columns was done at pre-determined time points. Water samples were collected for analysis of BAM concentration and enumeration of bacterial and protozoan cells. Sampling was done in an icebox (0 - 1 °C) to reduce bacterial growth and activity. At experimental termination, the columns were opened and filter material was collected from the inlet end, middle and outlet end of the columns for enumeration of bacterial cells and characterization of the prokaryotic community by molecular methods as well as enumeration of protozoan cells by most probable number (MPN) assay (see sections below). Finally, the BAM mineralisation potential of the cells attached to sand from the columns was confirmed by assessing the transformation of radioactively labelled BAM to ^14^CO_2_ in a standard mineralisation assay.

### Chemical analysis of 2,6-dichlorobenzamide

At each sampling point 15 ml of effluent water from each column was collected in 50 ml falcon tubes. To limit degradation during sampling, tubes were kept at 0 °C and the samples were stored at -18 °C until analysis. Storage time was maximum 5 months. At the day of analysis, samples were thawed and subsamples of 15 ml were extracted and pre-concentrated to 0.5 ml using solid phase extraction (SPE) as described in Albers et al. (2014). Prior to the SPE procedure, all samples including spiked calibration standards were spiked with 1 µg l^-1^ (final concentration after SPE) of deuterated BAM-d3 as an internal standard. BAM concentration was quantified using ultra performance liquid chromatography coupled to tandem mass spectrometry (ACQUITY UPLC system, Xevo TQ-S micro triple quadrupole; Waters Corporation). For additional method details see supplementary material (Table S1 and Table S2). LOD and LOQ for BAM was 2 ng l^-1^ and 6.8 ng l^-1^, respectively and recovery in effluent of the SPE was 96 %.

### Quantification of *bbdA* genes, MSH1 cells, and total 16S genes

Water samples of 100 ml were filtrated through a 0.2-μm MicroFunnel filter unit (Pall Corp., Ann Arbor, MI) immediately after extraction from the column. The filters were then transferred to bead tubes supplied with the DNeasy PowerWater Kit (Qiagen, København Ø, DK) and frozen for subsequent DNA extraction according to the manufacturer’s instructions. DNA was extracted from Nevtraco samples using the DNeasy PowerLyzer PowerSoil Kit (Qiagen, København Ø, DK) according to the manufacturer’s instructions.

Quantitative PCR targeting the *bbdA* gene responsible for the first step of BAM-degradation was performed in 20 µl reactions containing 1 µl DNA template, 4 µl HOT FIREPol® Evagreen® qPCR Supermix (Solis Biodyne), 4 pmol forward and reverse primers each. The primers bbdA-F (5’-ATATCACGGCCGGTACTATGCCAA-3’) and bbdA-R (5’-TCTTCCAAGATCGAACAACCCGGA-3’) (T’Syen et al. 2015) were used, amplifying a 156 bp PCR product including primers. Amplifications were done using the following running conditions: 95 °C for 12 min followed by 40 cycles of 98 °C for 10 s, 60 °C for 30 s, 72 °C for 2 min and finally 72 °C for 2 min followed by a melt curve prepared by increasing the temperature in 0.5 °C increments every 5 s from 60 to 95 °C.

A novel primer set was designed to verify that *bbdA* gene presence corresponded to presence of MSH1 cells. This primer set specifically targets a prophage-insertion-region on the genome of MSH1 (Nielsen et al. 2018), following an analogous method used by Gobbi et al. (2020). The prophage sequences in the MSH1-genome were identified by using PHASTER (PHAge Search Tool Enhanced Research) from Arndt et al. (2016). The chosen region was selected in order to partly cover a prophage sequence while including its insertion point within MSH1 genome, to increase the specificity toward our strain. Different primer sets were identified by using Primer-BLAST (Ye et al. 2012) and we selected a pair, which amplified a 91bp fragment, to increase qPCR efficiency due to the short amplicon size (Debode et al. 2017). The designed primers MSH1-F (5’-CATAGTTGGGCTGCGACAGG-3’) and MSH1-R (5’-CACTGGTTCTCACCATGGGC-3’) were applied in 20 µl reactions containing 1 µl DNA template, 4 µl HOT FIREPol® Evagreen® qPCR Supermix (Solis Biodyne), 4 pmol forward and reverse primers each. Amplifications were performed with the following running conditions: 95°C for 12 min followed by 40 cycles of 98 °C for 10 s, 55 °C for 30 s, 72 °C for 2 min and finally 72 °C for 2 min followed by a melt curve prepared by increasing the temperature in 0.5 °C increments every 5 s from 55 to 95 °C.

For both MSH1 numbers and *bbdA* genes standard curves were prepared from DNA extractions of a known number of MSH1 cells as determined by microscopical counting. Sample quantification was automatically done by Bio-Rad CFX manager 3.1.

Total bacterial numbers were determined by quantitative PCR targeting 16S rRNA gene performed according to Gobbi et al. (2019). In short, the primers used were 341F and 806R (Hansen et al. 2012), complete with adapters thus identical to those used for Illumina MiSeq sequencing. 16S rRNA gene standard curves were prepared from DNA extracts of *Escherichia coli* K-12 (Blattner et al. 1997, Feld et al. 2016) with 8.45 × 10^8^ 16S rRNA gene copies in the undiluted standard. *E. coli* K12 has seven 16S ribosomal RNA gene copies per genome, as compared to the average of four copies per bacterial genome (Větrovský and Baldrian 2013). 16S rRNA gene copy numbers were here automatically calculated from the 16S rRNA gene copies of the standard curve by Bio-Rad CFX manager 3.1. All qPCR analyses were performed on a CFX Connect(tm) Real-Time PCR Detection System (Bio-Rad, Hercules, CA).

### Enumeration of protozoa

Aliquots of water or Nevtraco samples were mixed with amoeba saline and then either thoroughly vortexed (water) or shaken for 30 min at 200 rpm (sand). The number of protozoa in the samples was estimated by a most probable number (MPN) assay in sterile 96-well microplates according to Ellegaard-Jensen et al. (2016). In short, dilution series were made and transferred to the microplate wells. Wells were inspected for the presence or absence of active protozoa at two pre-determined times (after one and three weeks) during incubation at 15 °C to ensure protozoan growth for visualisation.

### Library preparation and sequencing of the prokaryotic communities

DNA extracted from Nevtraco samples collected from the inlet end, middle and outlet end of the columns at the end of the experiment was used for amplicon library preparation for subsequent sequencing of the 16S rRNA gene. Library preparation was carried out by a two-step PCR as described earlier (Albers et al. 2018, Feld et al. 2016). In short, the V3-V4 region of the 16S rRNA gene was amplified using the 341F and 806R primers with adaptors. PCR reactions were conducted on a SimpliAmp Thermal Cycler (Applied Bio-systems, Foster City, California, US). In the first PCR, the PCR mixture contained 12 µl AccuPrime Supermix II (Invitrogen, Eugene, Oregon, US), 0.25 µM of each primer, 0.25 mg ml^-1^ BSA (Bioron), 5 µl template DNA (<0.05-2.5 ng DNA) and DNase free water to a final volume of 20 µl. The conditions of the first PCR were as described by Feld et al. (2016). Aliquots of the PCR products were run on a 1% agarose gel and after ethidium bromide staining checked under UV light. Following, the second PCR was done, to add indexes and sequencing adaptors, with 5 µl of product from the first PCR as template as described by Albers et al. (2018). The final PCR products were purified with 15 µl HighPrep PCR magnetic beads (MagBio Genomics Inc. Gaithersburg, Maryland, US) according to the manufacturer’s instructions and eluted in 27 µl TE buffer. Concentrations of the amplified and purified DNA samples were measured on a Qubit 2.0 fluorometer (Invitrogen, Eugene, Oregon, US). The samples were then equimolarly pooled, and this final library was sequenced on an Illumina MiSeq using the V2 kit (Illumina Inc. SanDiego, California, US) resulting in 2×250 bp reads.

### Bioinformatics analysis

Demultiplexed reads from Illumina sequencing were analysed using QIIME 2 v. 2018.11 (Bolyen et al. 2019, Hall and Beiko 2018). Reads were filtered, denoised, merged, chimera checked and dereplicated using the DADA2 R package v. 1.6.0 (Callahan et al. 2016) with default settings. Alignment and phylogenetic trees were generated using MAFFT (Katoh and Standley 2013) and FastTree (Price et al. 2010). Following inspection of rarefaction curves to check for saturation, the output was rarefied at 6.200 reads to include the sample with the lowest read number.

Taxonomic classification was performed using qiime feature-classifier in which a pre-trained Naïve-Bayes classifier with Silva v. 132 (silva-132-99-nb-classifier) was applied (Quast et al. 2013). Following, taxonomic tables were constructed through the qiime taxa barplot command incorporating the metadata, and the tables were exported for further processing.

The differences in the alpha diversity, based on number of OTUs, Faith’s phylogenetic diversity (Faith 1992) and Pielou’s evenness index were assessed by Kruskal-Wallis-Pairwise test. While the beta diversity of the column samples, based on Bray-Curtis dissimilarity measure, was visualised by PCoA plots through Emperor (Vázquez-Baeza et al. 2013) and analysed using PERMANOVA (Anderson 2001).

As a further analysis, sequences with ≥ 97% similarity to *Aminobacter* sp. MSH1 were removed from the dataset, to assess effects of the treatments on the indigenous bacterial community of the water. Here samples were rarefied at 1.550 reads as a consequence of filtering out a high number of the reads (i.e. those with ≥ 97% similarity to *Aminobacter* sp. MSH1). All other steps of the analysis were done as described above.

Stacked bar charts were created in SigmaPlot (version 13.0) and heatmap of the most dominant indigenous prokaryotic OTUs was made in R (www.rproject.org).

### Statistics

Statistical analyses, besides the bioinformatics described above, were performed as t-tests in SigmaPlot v. 13.0 (Systat Software Inc). Differences between treatments were found to be statistically significant when p ≤ 0.05. Unless otherwise stated, data are presented as mean ± standard error (SE).

### Data availability

The sequence data have been deposited and are publicly available from NCBI database (BioProject ID: PRJNA602585).

## RESULTS

### Prolonged performance of the sand filters

BAM concentrations in the outlet from the inoculated columns were below detection limit throughout the entire experiment. This represents the complete removal of the spiked 3.0 µg l^-1^ BAM in the membrane retentate water and of the 0.3 µg l^-1^ BAM in the untreated inlet water. The two non-inoculated control columns did initially not show any removal of BAM, however a gradual decline in BAM concentration in the outlet water was detected for the control column receiving membrane retentate water ending with 1.4 µg l^-1^ BAM left in the outlet water (Fig. 1).

**Figure 1.**
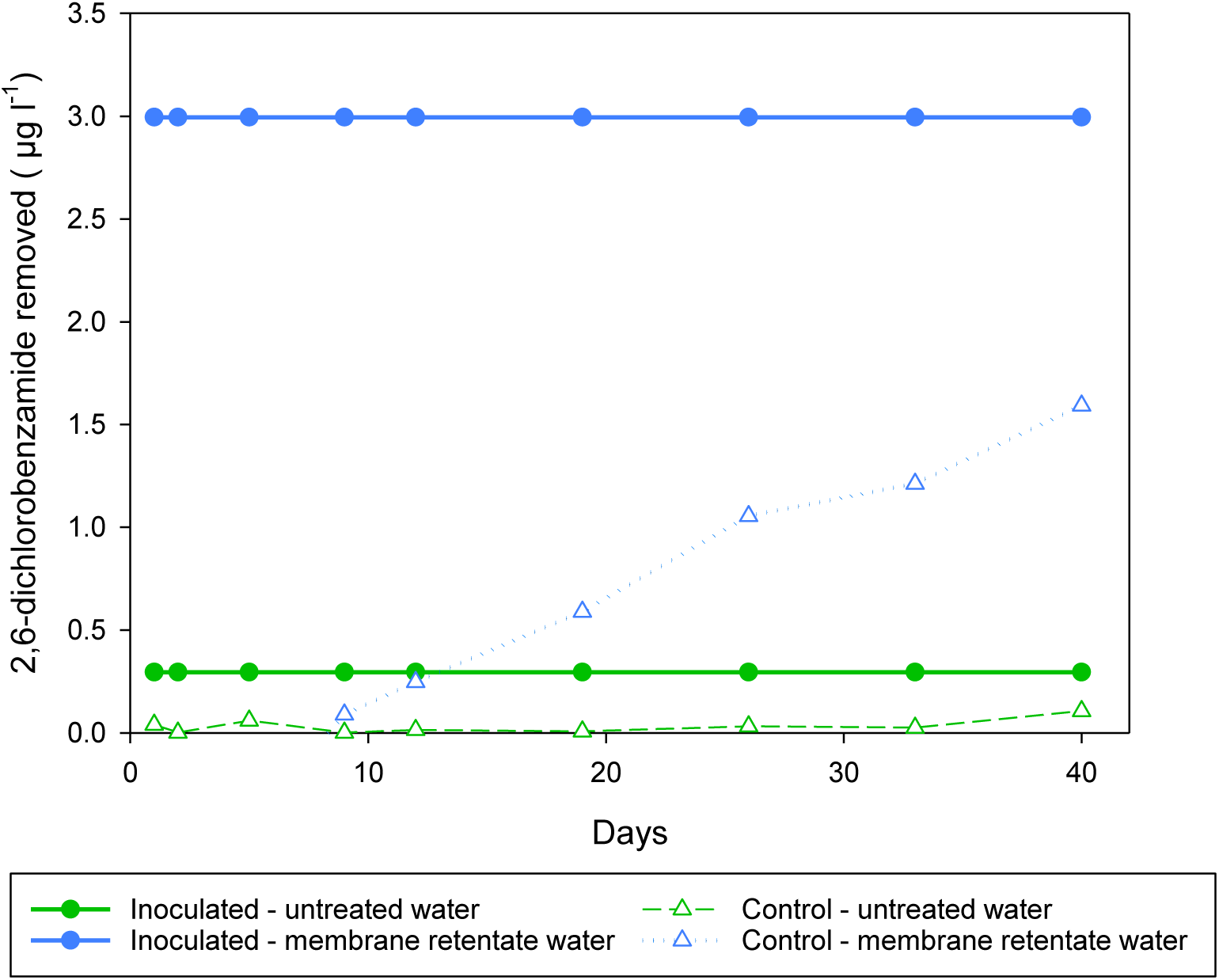
Amount of 2,6-dichlorobenzamide (BAM) removed in the columns. Circles represent columns inoculated with BAM-degrader *Aminobacter* sp. MSH1, while triangles represent control columns. Columns received as feed either untreated water with 0.3 µg l^-1^ BAM (green) or membrane retentate water with 3.0 µg l^-1^ BAM (blue).

### Survival and persistence of *Aminobacter* sp. MSH1

The survival and persistence of *Aminobacter* sp. MSH1 in the columns were measured as its presence in the outlet water determined by qPCR targeting the specific degrader gene *bbdA* at selected time points. The highest number of MSH1 cells in the outlet of the columns was detected at the start of the experiment, i.e. following the initial ∼22 hours after inoculation and initiation of flow with MSNC medium as inlet, with on average 1.6 × 10^7^ cells ml^-1^ for all the inoculated columns. Equal to an average loss of 31 % (range 20 - 57 %) of all cells initially inoculated into the columns. There was no significant difference found (p = 0.191) between the amount of MSH1 cells in the outlet of the inoculated columns which would subsequent be fed with membrane retentate or untreated water (Fig. 2). Following the switch of feed from MSNC media to untreated or membrane retentate water with BAM, cell loss dropped to approximately 4 × 10^5^ cells ml^-1^ for both treatments in the samples taken on the day immediately after the switch of feed equalling a loss of less than 1 % of the total amount of MSH1 cells added to each column. At the following sampling points, the cell losses were even lower and from day 19 most values were below quantification limit (Fig. 2).

**Figure 2.**
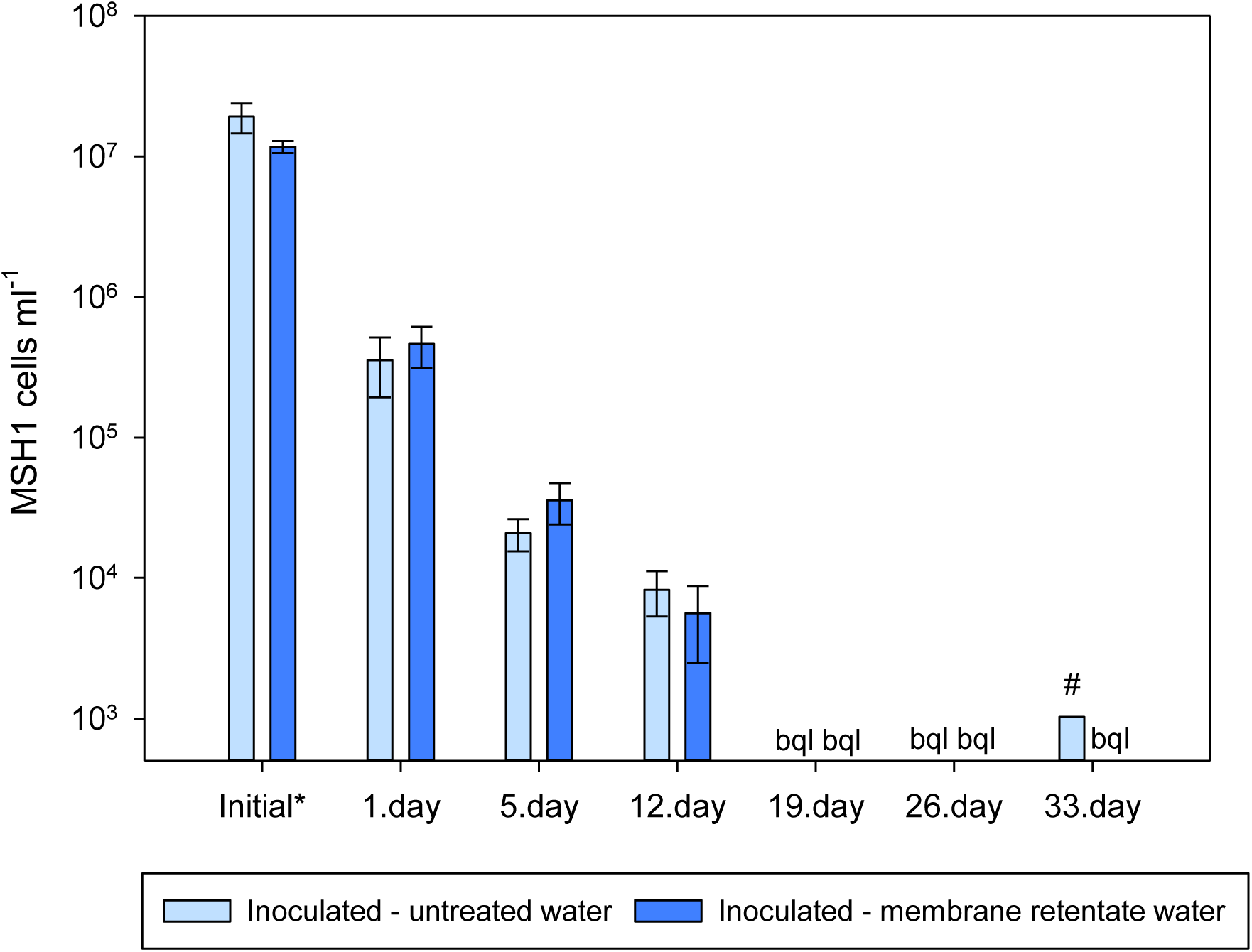
Number of *Aminobacter* sp. MSH1 cells detected in the outlet of the inoculated columns based on the quantification of *bbdA* genes. *Indicates the initial loss of MSH1 which was sampled during the first 22h of operation after inoculation and inlet flow of MSNC was started. The rest of the sample times (1-33 days) indicate time after the columns first received untreated (light blue bars) or membrane retentate water (dark blue bars) with BAM. bql: below quantification limit. # only one sample out of three was above quantification limit. Data presented are means with error bars showing standard error.

At experimental termination, the number of MSH1 attached to the Nevtraco material was determined by qPCR for material collected from the inlet end, the middle and outlet end of the columns. This showed a large number of MSH1 persisting throughout the entire column with numbers decreasing slightly from the inlet end towards the outlet end of the column for all the inoculated columns (Fig. 3). There was no significant difference between the number of MSH1 attached to the sand in the columns when comparing those receiving membrane retentate and untreated water (inlet, p = 0.238; middle, p = 0. 239; outlet, p = 0.140).

**Figure 3.**
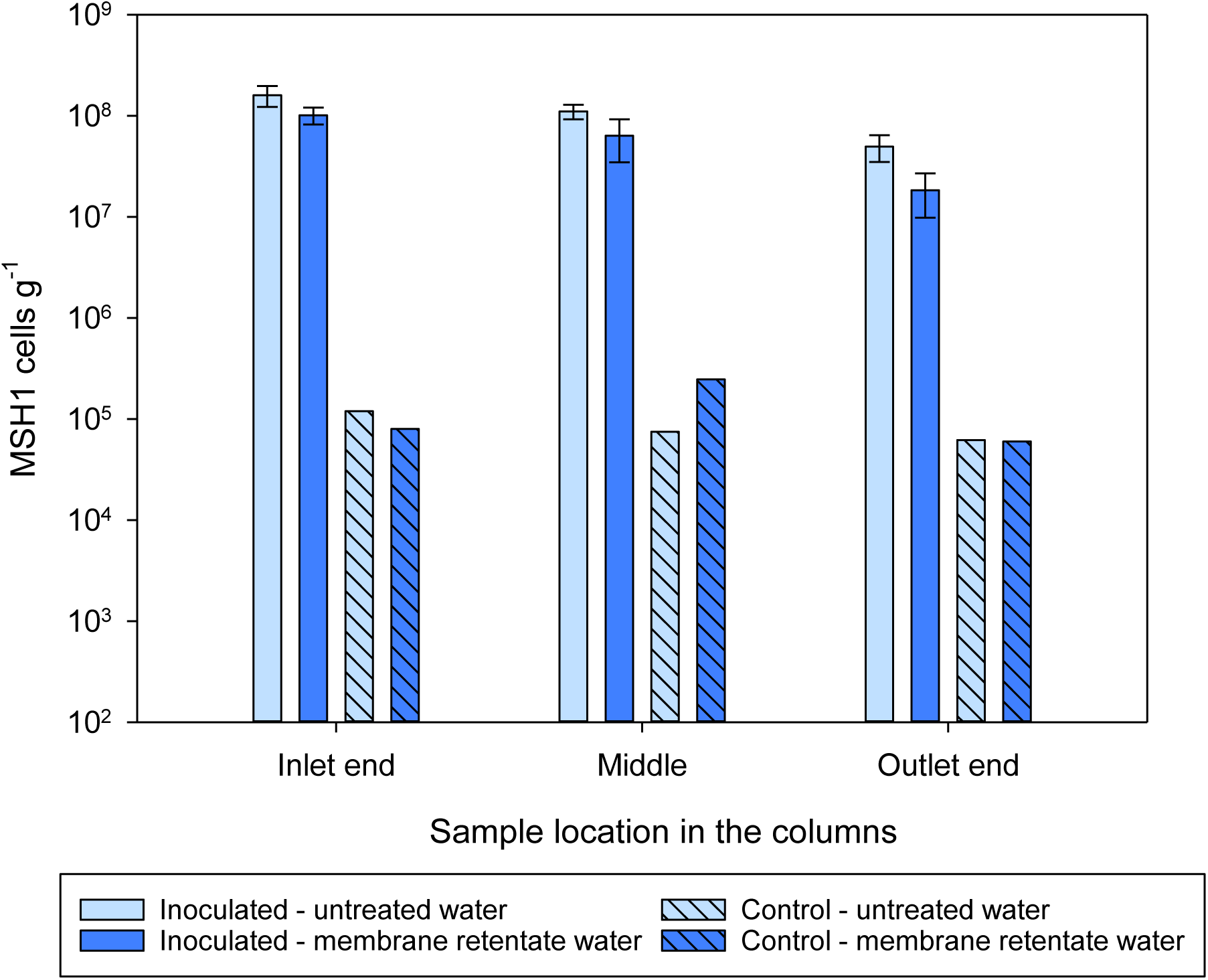
Number of *Aminobacter* sp. MSH1 cells based on the quantification of *bbdA* genes detected in the sand sampled from different locations within the columns at experimental termination. Columns received during operation a feed of either untreated water with 0.3 µg l^-1^ BAM (light blue bars) or membrane retentate water with 3.0 µg l^-1^ BAM (dark blue bars). For the columns inoculated with *Aminobacter* sp. MSH1data are presented as means (n = 3) with error bars showing standard error. Two control columns were run without initial inoculation (hatched bars; n = 1).

The *bbdA* gene was also detected in the material collected from the non-inoculated control columns. However, the number of MSH1 cells in the control columns were in the range of 10^4^ - 10^5^ cells g^-1^ – in contrast to the 10^7^ - 10^8^ MSH1 cells g^-1^ determined in the inoculated columns – and in all sample locations within the columns below 0.5 % of the amount determined in the inoculated columns at the same respective sample locations (Fig. 3).

These findings were verified by the number of cells detected by qPCR applying the newly designed MSH1 specific primers (Fig. S1). Thus confirming that presence of the *bbdA* gene corresponded well with the number of MSH1 cells determined by this specific primer set. Further, this confirms that 1) the *bbdA* gene was not lost by MSH1 in our experiment, and 2) no other organism harboured the *bbdA* gene and was thus responsible for the BAM degradation.

### Protozoan abundance

As the columns received feed harbouring its natural water microbial community, and in the case of the membrane retentate containing natural microbes in even higher densities (See table 1), inoculated MSH1 cells experienced predator-prey interactions from this microbial community. We here investigated the number of grazers (i.e. protozoa) in the water at selected time points and on the filter material as well as total bacterial numbers and the prokaryotic community compositions in the columns.

While the membrane retentate water contained approximately 10 - 600 protozoa ml^-1^, the untreated feed water had much lower numbers ranging from below the quantification limit to approximately seven protozoa ml^-1^. Protozoan numbers in the outlet water from the columns were similar (i.e. not significantly different; p = 0.929) for the inoculated columns of both treatments with approximately 120 protozoa ml^-1^ nearly half way through the experiment (day 16), while in the outlet of the control numbers did not exceed four protozoa ml^-1^ at this point in time (Fig. 4A). Near the end of the experiment (day 36), the number of protozoa in the outlet of the inoculated columns receiving retentate water was slightly higher than the number from those receiving untreated water (Fig. 4A), though the difference was not statistically significant (p = 0.117).

**Figure 4.**
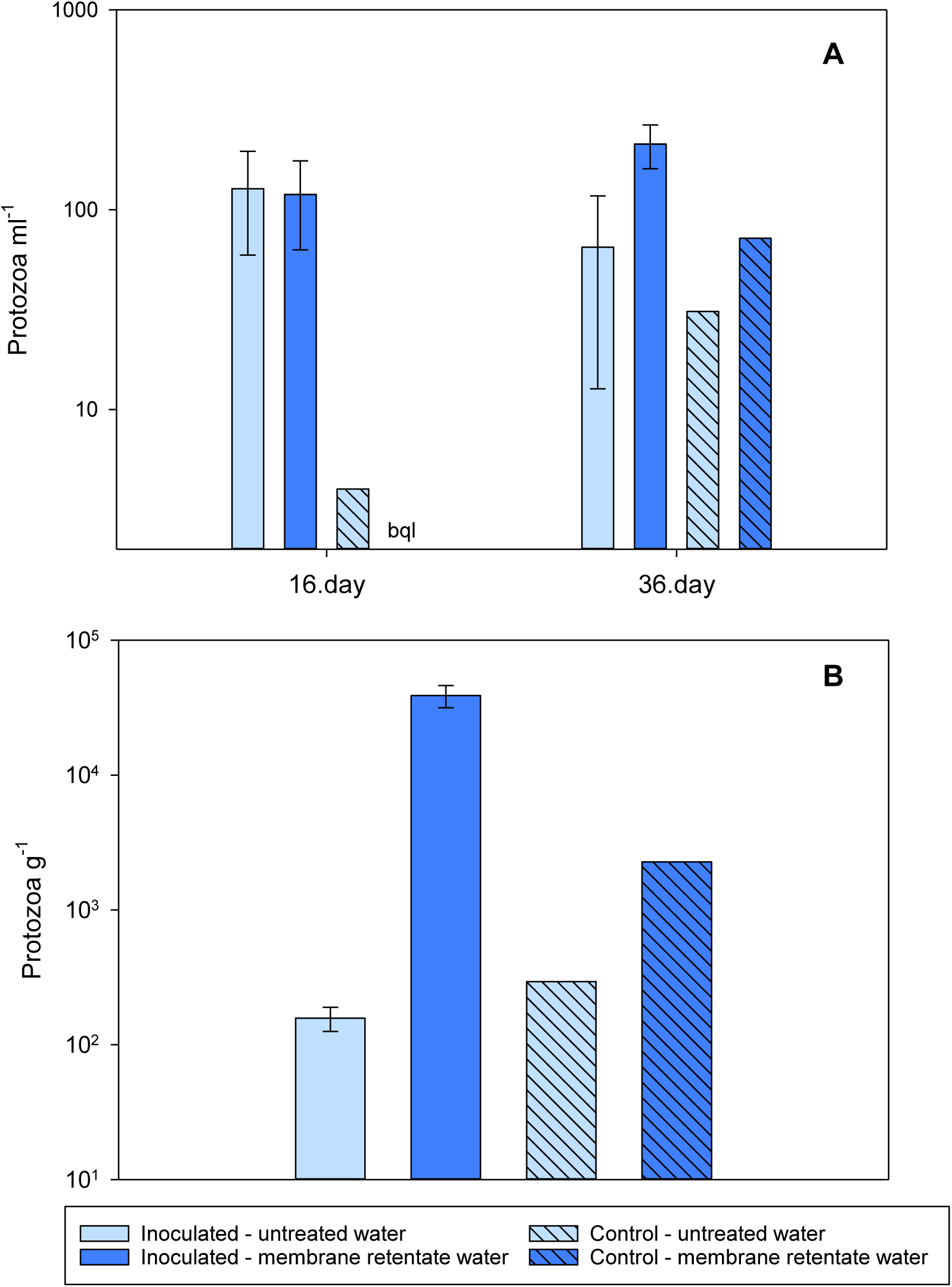
Number of culturable protozoa in A) the outlet of the columns and B) the sand of the columns at experimental termination. Columns received during operation a feed of either untreated water with 0.3 µg l^-1^ BAM (light blue bars) or membrane retentate water with 3.0 µg l^-1^ BAM (dark blue bars). For the columns inoculated with *Aminobacter* sp. MSH1 data are presented as means (n = 3) with error bars showing standard error. Two control columns were run without initial inoculation (hatched bars; n = 1). bql: below quantification limit.

At the end of the experiment, the number of protozoa attached to the Nevtraco material in the inlet end of the columns was 200× higher for the inoculated columns receiving membrane retentate water than those receiving untreated water (p = 0.006). While for the number of protozoa in the control columns, approximately 10× more protozoa was found in the column receiving membrane retentate water compared to the column receiving untreated water (Fig. 4).

### Total bacterial communities in the columns

The total number of bacteria on the filter material in the columns was determined by qPCR of the 16S rRNA gene. The pattern for 16S rRNA genes in the inoculated columns reflected that of the *bbdA* gene (Fig. 3 and 5) with numbers decreasing from the inlet towards the outlet end. A significant difference for the number of 16S rRNA genes in the inoculated columns receiving untreated and membrane retentate water was only found for the outlet end of the columns (inlet, p = 0.120; middle, p = 0.198; outlet, p = 0.045). Total bacterial numbers in the control columns also decreased from the inlet towards the outlet end, ending with values below quantification limit, and the numbers were approximately 15× and 70× lower for untreated and membrane retentate water columns, respectively, compared to the inoculated columns.

**Figure 5.**
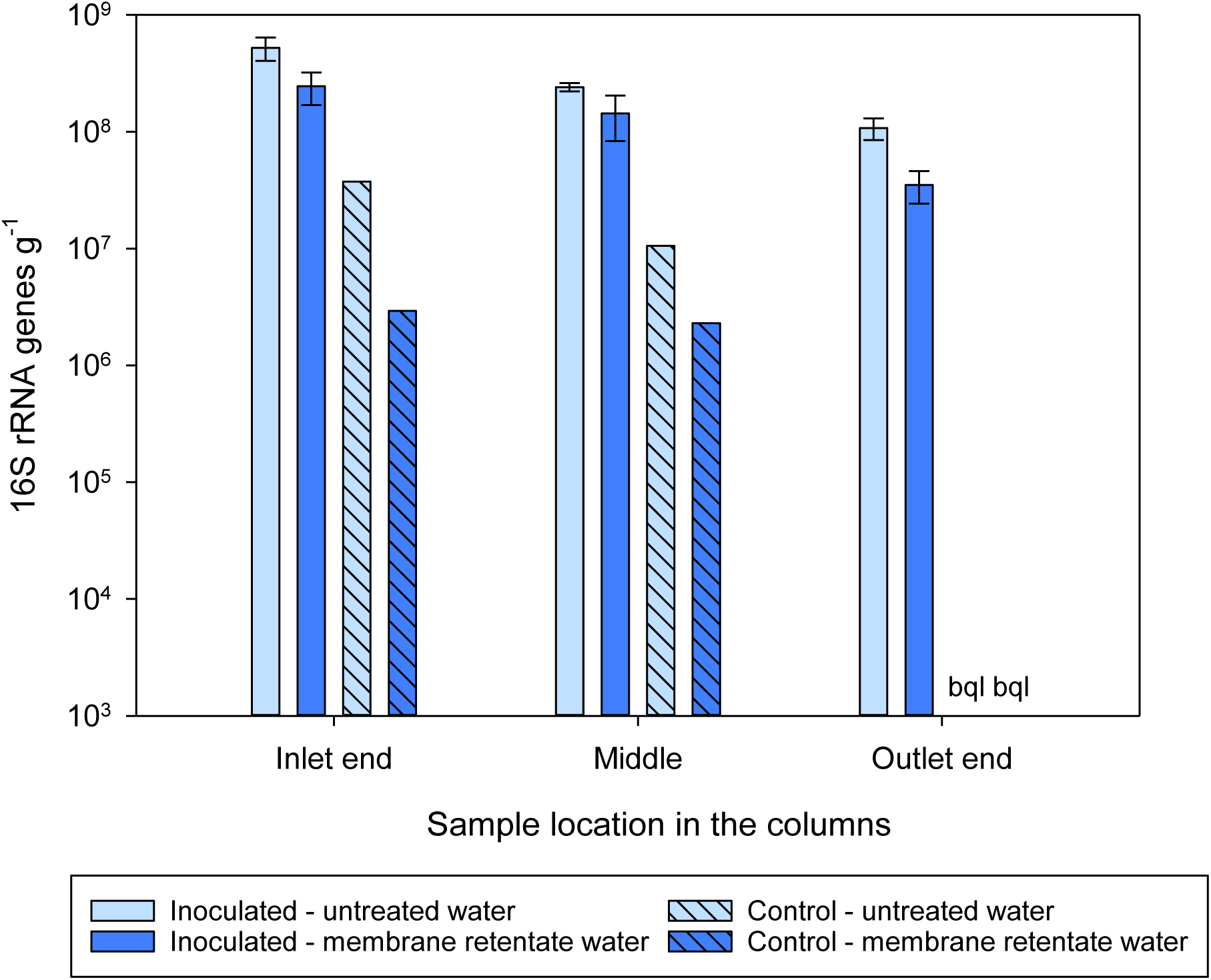
Number of total bacteria determined by detection of the 16S rRNA gene on the sand sampled from different locations within the columns at experimental termination. Columns received during operation a feed of either untreated water with 0.3 µg l^-1^ BAM (light blue bars) or membrane retentate water with 3.0 µg l^-1^ BAM (dark blue bars). For the columns inoculated with *Aminobacter* sp. MSH1 data are presented as means (n = 3) with error bars showing standard error. Two control columns were run without initial inoculation (hatched bars; n = 1). bql: below quantification limit.

Results from the amplicon sequencing showed that prokaryotic communities in all the columns were dominated by Proteobacteria (Fig. 6A). The inoculated columns were dominated by Alphaprotobacteria with a relative abundance of 86.9%, 95.6% and 85.3% in the inlet, middle and outlet end, respectively, in the columns receiving untreated water and 92.0%, 90.0% and 78.4% in the inlet, middle and outlet end, respectively, in the columns receiving membrane retentate water. This reflects the dominant presence of MSH1 since the 16S phylogenetic origin of *Aminobacter* sp. MHS1 lies within Alphaprotobacteria in the *Rhizobiaceae* family, which we will go more into detail with in the sections below. In the control columns the relative abundance of Alphaprotobacteria ranged from 24.3 % - 49.4%, while a higher relative abundance of Gammaproteobacteria (37.4% - 56.8%) was seen in those columns compared to the inoculated columns.

**Figure 6.**
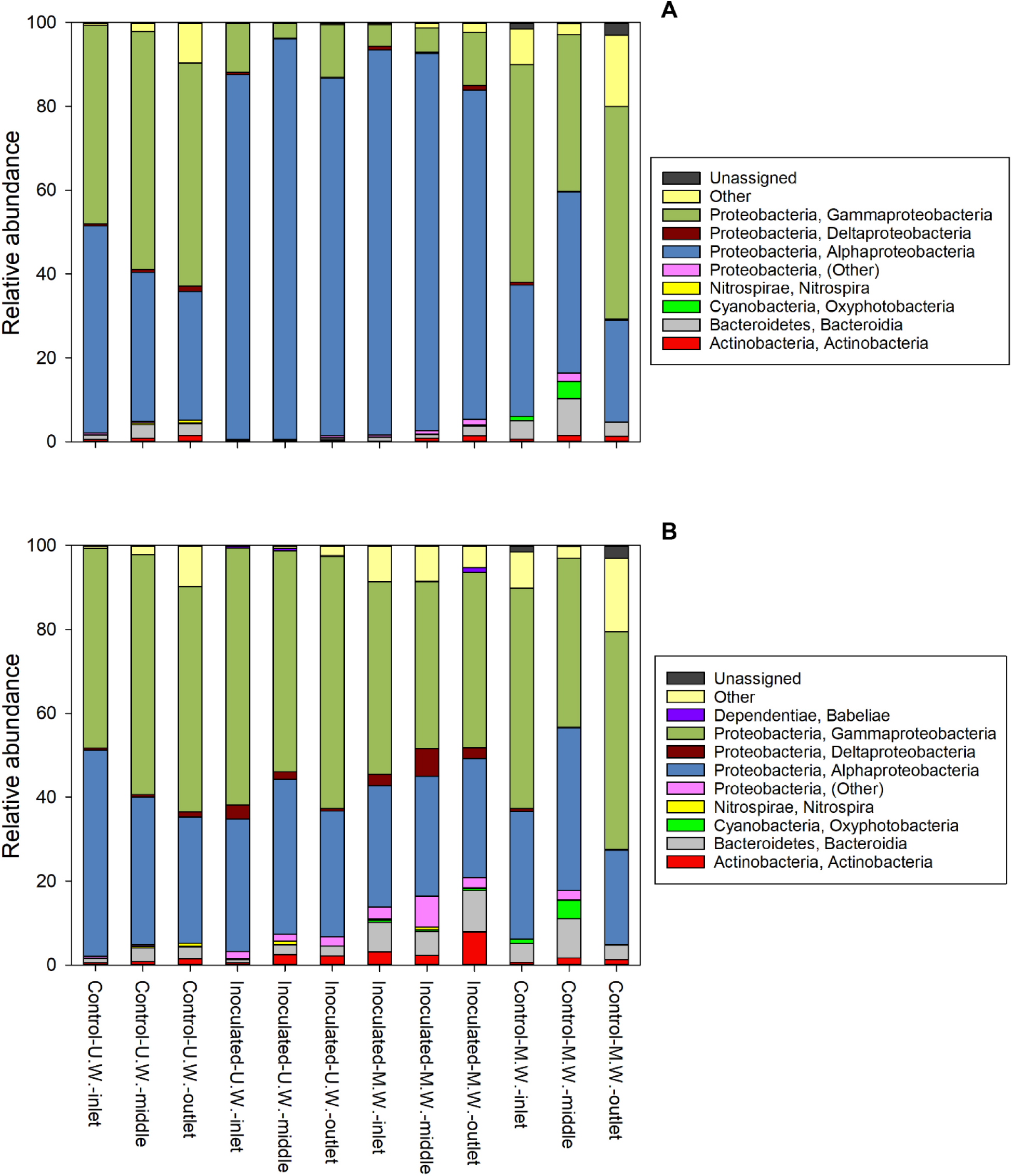
Composition of the prokaryotic communities of the columns at day 40. A) the relative abundances within the whole communities and B) the relative abundances within the dataset where sequences with ≥ 97% similarity to *Aminobacter* sp. MSH1 had been removed. The boxes to the right show at class level the identities of the 16S rRNA genes that had an accumulated relative abundance of >1% across all samples. M.W.: Columns receiving membrane retentate water. U.W.: Columns receiving untreated water.

Investigating the microbial communities of the inoculated columns more closely at order level, we found that *Rhizobiales* is dominating with a relative abundance of 90.3%, 88.2% and 73.5% in the inlet, middle and outlet end, respectively, in the columns receiving membrane retentate water and 83.2%, 94.1% and 80.9% in the inlet, middle and outlet end, respectively, in the columns receiving untreated water. In contrast, we found only 13.3%, 15.0% and 2.9% *Rhizobiales* in the inlet, middle and outlet end, respectively, of the control columns receiving membrane retentate water, and approximately 2% in all three locations in the control columns receiving untreated water. Generally, the control columns showed a higher diversity for all three alpha diversity measures (number of OTUs, Faith’s Phylogenetic Diversity, and Pielou’s evenness) compared to the inoculated columns (Table S3) reflecting that the diversity of the inoculated columns was reduced by a dominating organism. However, the inoculated columns receiving the membrane retentate water showed to have a higher diversity compared to the ones receiving untreated water (6.5 and 5.8 Faith’s Phylogenetic Diversity, respectively).

Since *Rhizobiales* is the order *Aminobacter* belongs to, we examined this taxonomic group more closely. Focusing on members of this group in the OTU table, we found 15 different OTUs belonging to the *Rhizobiales* order, but one stood out with a relative abundance of ∼ 80% of the total prokaryotic communities in the inoculated columns. This OTU belong to the *Rhizobiaceae* family, but was not further classified. Exploring the 67 features (unique sequence variants) of this OTU, we found that one had a 100% match with *Aminobacter* sp. MSH1 when conducting a BLAST (Basic Local Alignment Search Tool) search in NCBI (https://blast.ncbi.nlm.nih.gov/Blast.cgi). This feature was the overall most frequent constituting 66.8% of all the quality checked and filtered sequences of the total dataset underlining the dominance of *Aminobacter* sp. MSH1 in the systems. The relative abundances of *Aminobacter* sp. MSH1 in the columns support this further with values ranging 70.9 – 93.1 % of the total prokaryotic communities in the inoculated columns (Fig. S2). In contrast, the second and third most frequent features constituted 4.5% and 3.2%, respectively, and belonged to the *Pseudomonas* and *Sphingobium* genera.

To investigate the effect that *Aminobacter* sp. MSH1 had on the rest of the community, all sequences were re-analysed filtering out sequence variants that had ≥ 97 % similarity to MSH1. At class level this resulted in a higher similarity of the relative abundances of the prokaryotic communities of the columns (Fig. 6B), with Gammaproteobacteria as the most abundant group on average. This also revealed the class *Babeliae* in the phylum *Dependentiae*, which are often found to be pathogens of diverse aquatic protists (Deeg et al. 2019), to be present throughout the inoculated columns and in the inlet end of the control column receiving untreated water (Fig. 6B) in up to 1.3% relative abundance. On the other hand, when inspecting the heatmap of the most abundant OTU’s, just a few are only present in the control columns; *Rhodoferax* sp. and *Rickettsiaceae* sp., but only in the column receiving untreated water (Fig. S3). When inspecting the beta diversity of the communities, after filtering out the MSH1 sequences, they very distinctly grouped according to the feed that they received i.e. treated and untreated water (Permanova p = 0.01; Fig. 7).

**Figure 7.**
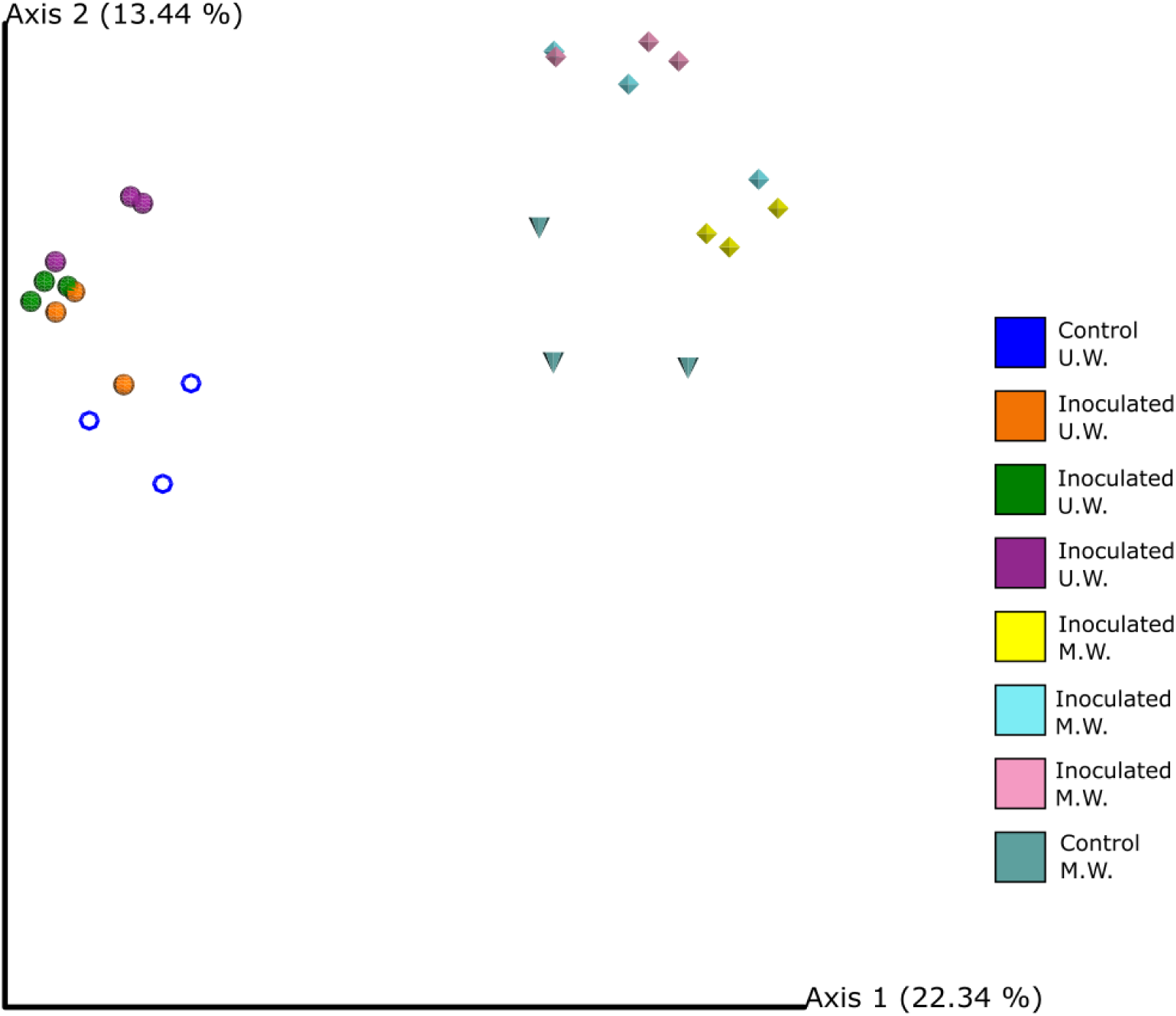
The beta diversity of the prokaryotic communities of the columns at day 40 where sequences with ≥ 97% similarity to *Aminobacter* sp. MSH1 had been removed from the dataset. The PCoA shows Bray-Curtis dissimilarity matrix calculated through Emperor. Columns received during operation a feed of either untreated water with 0.3 µg l^-1^ BAM (U.W.) or membrane retentate water with 3.0 µg l^-1^ BAM (M.W.). Individual colours represent samples from the same column. The two control columns run without initial inoculation are represented by the blue circles and blue cones, respectively.

## DISCUSSION

This study provides the first evidence that bioaugmented sand filters can be used for efficient treatment of membrane retentate water in a drinking water treatment system, as we here show a 100% removal of the pesticide residue BAM by bioaugmented sand filtration. This removal efficiency is superior to that of previously published studies, which found as a maximum around 65 – 80 % BAM removal in bioaugmented sand filtration (Albers et al. 2015b, Ellegaard-Jensen et al. 2016, Horemans et al. 2016). Furthermore, as opposed to field studies that found that the degradation efficiency declined over time (Albers et al. 2015b, Horemans et al. 2016), we were able to maintain the same high removal efficiency throughout the 40 days long experiment.

Unexpectedly, we found the same high removal efficiency in both sand columns receiving membrane retentate water as well as those receiving untreated water. A result which was supported by a comparable number of MSH1 cells in the two types of columns, and showing that we were successful in prolonging a sufficient survival, persistence and activity of the MSH1 cells added to the sand filter columns. Two possible reasons for this, which will be discussed in detail below, are: 1) Our newly developed inoculation strategy enabled the added bacteria to achieve a stronger attachment to the sand through biofilm formation resulting in less washout of the added cells, as well as providing a competitive and protective advantage against the indigenous microbes of the water, and 2) that flow rate (similar in both types of columns, but lower than previously reported experiments) is decisive in the washout of cells and thus governs obtained degradation efficiency.

The inoculation strategy developed and applied in our experiment added a step where a growth medium was applied as feed to the columns first 48 hours following the inoculation. This should in theory allow the added cells to grow and form a biofilm on the sand grains. The initial loss, during the first 22 hours, of 31 % (range 20 - 57 %) cells of the total amount of MSH1 cells added to each column corresponds well with previous studies. In those the initial loss of inoculated MSH1 cells were 20 - 50 % during the first two to three hours, followed by a loss of 1 - 27 % during the following approximately 24 hours (Albers et al. 2015b, Albers et al. 2014, Ellegaard-Jensen et al. 2016). However, growth of the population added must be assumed, in the present study, given the optimal growth medium supplied. Indeed Schultz-Jensen et al. (2014) reported a growth rate for MSH1 of μ = 0.04 h^−1^ at 10 °C in a batch system. While Sekhar et al. (2016) reported growth and biofilm formation by MSH1 at concentrations as low as 1 µg l^-1^ BAM in flow channel systems, with an increasing biofilm thickness at higher substrate concentrations. Indeed, we found that a large population of MSH1 was sustained in the sand filter columns throughout the present study despite the protozoan presence, enabling complete BAM removal as opposed to previous experiments (Albers et al. 2015b, Ellegaard-Jensen et al. 2016, Horemans et al. 2016) as mentioned above.

Horemans et al. (2017) found that MSH1 was able to successfully invade a sand filter community biofilm and achieve maximum 60% BAM removal under oligotrophic conditions. While the possible protective mechanism of our approach of allowing MSH1 to first form biofilm on the sand before exposing it to the indigenous microbiota remains to be determined through further studies, we provide some insight into possible interactions between the inoculated strain and the indigenous bacterial community colonizing the sand through sequencing of the prokaryotic biofilm community.

The bacterial communities of the inoculated columns were as expected dominated by MSH1. However, a large number of other bacteria also colonised the sand harbouring roughly 50 - 150 OTUs. When sequences belonging to MSH1 where filtered out, patterns in the communities of the different columns were detected. Interestingly, this showed that the class *Babeliae* in the phylum *Dependentiae* was present throughout the inoculated columns and in the inlet end of the control columns (Fig. 6B). Members of the phylum *Dependentiae* (previously called TM6) are often found to be pathogens of diverse aquatic protozoa (Deeg et al. 2019, Delafont et al. 2015). Though found to be only a small part of the total bacterial community. This is in line with the high number of protozoa found especially in outlet from the inoculated columns, though the protozoan enumeration was based on culture dependent techniques, so a direct comparison cannot be made. *Legionella* sp. is another amoebal endosymbiont found in higher relative abundance in the inoculated columns receiving membrane retentate water compared to the other columns (Fig. S3). The natural habitat of *Legionella* is the aquatic environment including ground- and drinking water (Costa et al. 2005, Riffard et al. 2001, Wullings et al. 2011). A BLAST search (including only cultured organisms) in NCBI of the most dominant *Legionella* sequence variant resulted in a > 98% similarity to *Legionella* sp. leg101 recovered from DWTP by amoebal co-culture method (Corsaro et al. 2010). Though Corsaro et al. (2010) reported no *L. pneumophila*, other *Legionella* sp. may also be human pathogens (Gomez-Valero et al. 2014). Thus the increase in the relative abundance of *Legionella* sp. seen in our study warrants a deeper investigation into inoculation strategy and feed impact on prevalence of human pathogens, which is out of the scope of the present study. We suggest that strain specific methods are needed for this investigation, and if such investigations confirm pathogen proliferation, bactericidal treatment i.e. UV-treatment may be required if the purified water is to be utilized for drinking water.

Opposite to the above, an OTU belonging to the family *Rickettsiaceae* was only found in the control columns receiving untreated water (Fig. S3). The *Rickettsiales* are endosymbionts of diverse eukaryotic cells including that of amoebae in drinking water networks (Delafont et al. 2016). We speculate that the membrane treatment may led to rupture of some amoebae within the water, though this remains to be confirmed by further study. Delafont et al. (2014) further showed an amoebae–mycobacteria association in the drinking water network of Paris. However, no mycobacteria were detected in our columns, which suggests that the indigenous microbial communities are governed by the source of the water as well as the treatment conditions of the DWTP e.g. the age of the sand filter (Albers et al. 2015a, Bugge Harder et al. 2019).

Finally, the persistence of the inoculated cells in the present study is likely also affected by the flow rate of the columns. The columns had an approximate water residence time of 3 h, which is three to six times slower than that of Horemans et al. (2016) and approximately 12 times slower than that of Albers et al. (2015b). However, ten times longer residence time in the sand filters is also made possible through the suggested membrane filtration technology, where only a fraction of the water stream i.e. the membrane retentate is to be treated by sand filtration. Nevertheless, the effect of residence time on the persistence and performance of *Aminobacter* sp. MSH1 in sand filters remains to be established under controlled conditions.

Noticeably the flowrate and conditions of inoculation applied in the present study gives a superior persistence of the inoculated strain and BAM removal efficiency compared to previous studies as mentioned above. This is true even in the face of a higher grazing pressure (Fig. 4A and 4B) and a higher diversity of indigenous bacteria (Table S3) in the column receiving the membrane retentate water. That the indigenous prokaryotic community is affected by the column feed (Fig. 7) can be ascribed to the composition of elements in the untreated and retentate water (Table 1). Here especially non-volatile organic carbon (NVOC) will provide a foundation for increased microbial growth, and depending on the composition of the NVOC also a shift in the microbial community of the membrane retentate water compared to the untreated water. We suggest that further studies characterize the NVOC of membrane retentate in drinking water treatment, specifically focusing on the assimilable organic carbon (AOC).

In this study we successfully achieved full removal of pesticide residue in bioaugmented sand filter columns. We further show that membrane retentate should not be discarded as a waste product, but may be used as a valuable source of nutrients and applied as feed for bioaugmented sand filter columns.

## ACKNOWLEDGEMENTS

This project was funded by MEM2BIO (Innovation Fund Denmark, contract number 5157-00004B) and by Aarhus University Research Foundation starting grant (AUFF-E-2017-7-21). TREFOR Vand, DIN Forsyning A/S and HOFOR A/S are thanked for kindly providing access to drinking water treatment plants. Silhorko-Eurowater is thanked for useful discussion and for providing Nevtraco material.

## Supplementary material

### BAM concentration measured using ultra performance liquid chromatography–tandem mass spectrometry

#### Chromatography

Chromatography was done using an Acquity UPLC HSS T3 C18, (100 mm 2.1 mm, 1.8 µm) on a ACQUITY UPLC system (Waters Corporation). Column temperature was set at 40 °C during analysis. A constant flow rate of 0.5 ml/min was used. The mobile phase composed of solvent A (Ultra pure MilliQ water with 0.05% Formic acid) and solvent B (100% LC-MS grade acetonitrile). The gradient program is seen in table S1. The injection volume was 10 µl.

**Table S1:**
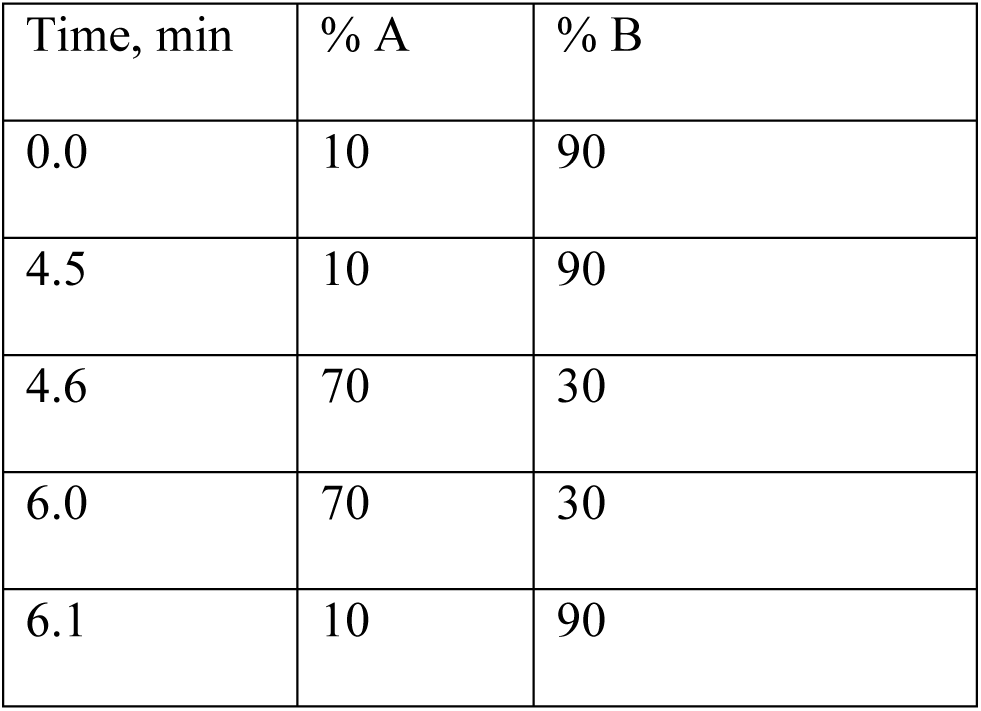
UPLC gradient program (8 min total run time)

| Time, min | % A | % B |
| --- | --- | --- |
| 0.0 | 10 | 90 |
| 4.5 | 10 | 90 |
| 4.6 | 70 | 30 |
| 6.0 | 70 | 30 |
| 6.1 | 10 | 90 |

#### Mass spectrometry

Detection of BAM was done on a Xevo TQ-S micro triple quadrupole (Waters) using positive electrospray ionization and capillary voltage of 2 kV. The desolvation and source temperature was set to 600 and 150 °C, respectively. Nitrogen was used as cone and desolvation gas with a flow of 30 and 1000 l/h, respectively. Argon was used as collision gas with at a pressure of 3.57 × 10^−3^ mBar.

BAM was identified using multiple-reaction monitoring (MRM) transitions with the settings shown in Table S2. To circumvent crosstalk of BAM-d3 in the BAM ion transitions, the Cl^37^ peak of BAM-d3 was used in order to increase the differences in m/z of the two parent ions.

**Table S2.** Retention time, MRM settings and operating parameters for the analysis of BAM and deuterated BAM-d3. MRM transitions are listed with the quantifier transition on top and the qualifier transitions below.

| Compound | Retention time | MRM transitions<br>m/z | Cone voltage, V | Collision<br>energies, eV |
| --- | --- | --- | --- | --- |
| BAM | 4.2 | 189.95>172.90 | 30 | 17 |
|  |  | 189.95>108.90 |  | 36 |
|  |  | 189.95>145.00 |  | 26 |
| BAM-d3 | 4.1 | 194.95>112.95 | 30 | 37 |
|  |  | 194.95>149.99 |  | 26 |
|  |  | 194.95>177.90 |  | 16 |

Data analysis with automated data processing was done using the Target Lynx software in MassLynx (Water corporation). Quantification of BAM was performed by integration of the area under the curve from the MRM chromatogram m/z 189.95>172.90 and m/z 193.00>112.95 of the BAM and the internal standard BAM-d3, respectively. The response (the ratio of the integrated area of BAM and BAM-d3) was used for quantification and the concentration of BAM was calculated from a calibration curve of spiked samples using linear regression. The calibration curve covered the concentration range of 0.005-1.5 µg/l BAM and was linear in the full range.

### Prokaryotic communities in the sand filter columns

**Table S3.**
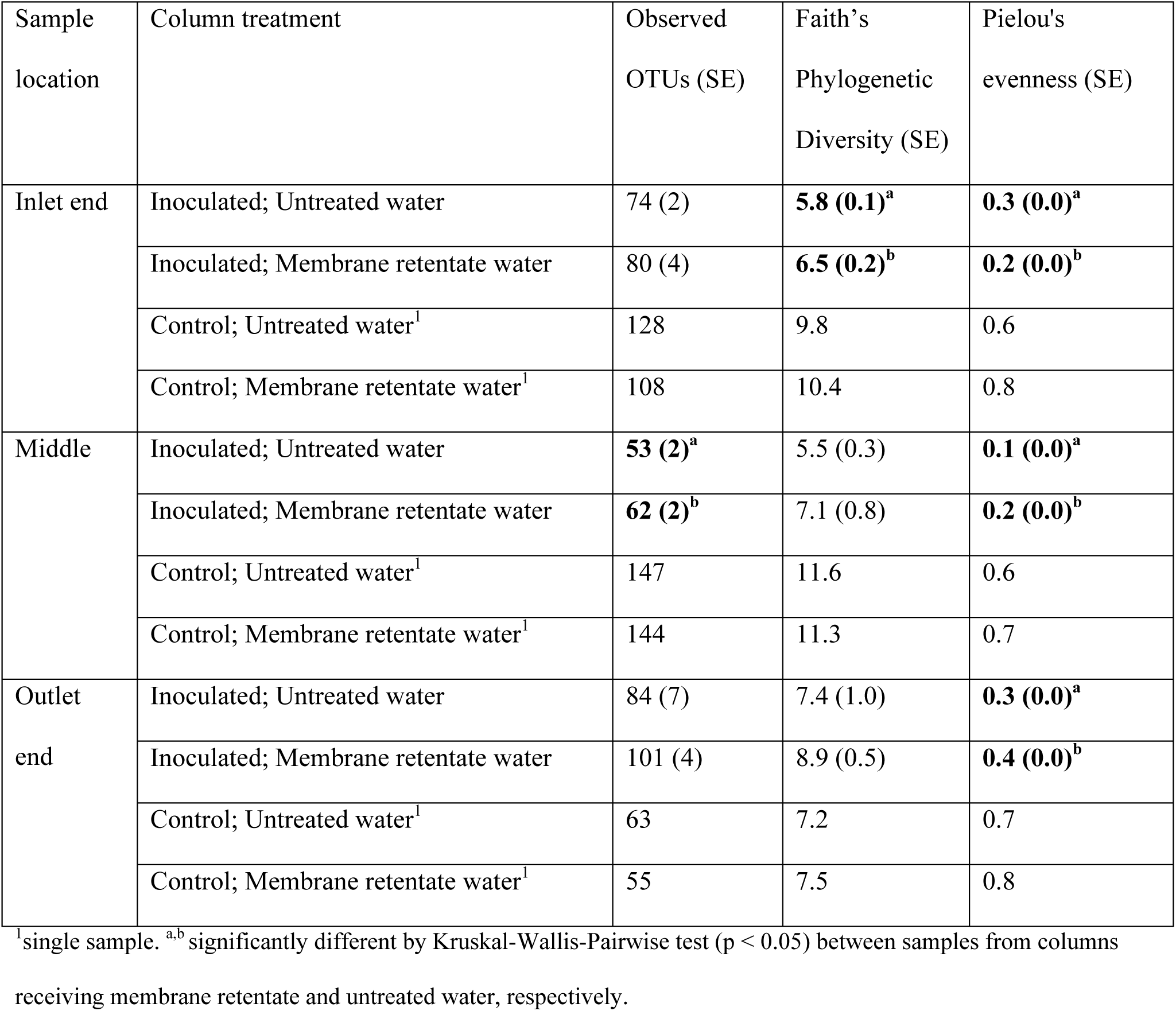
Alpha diversity of the total prokaryotic communities within the sand filter columns

**Fig. S1:**
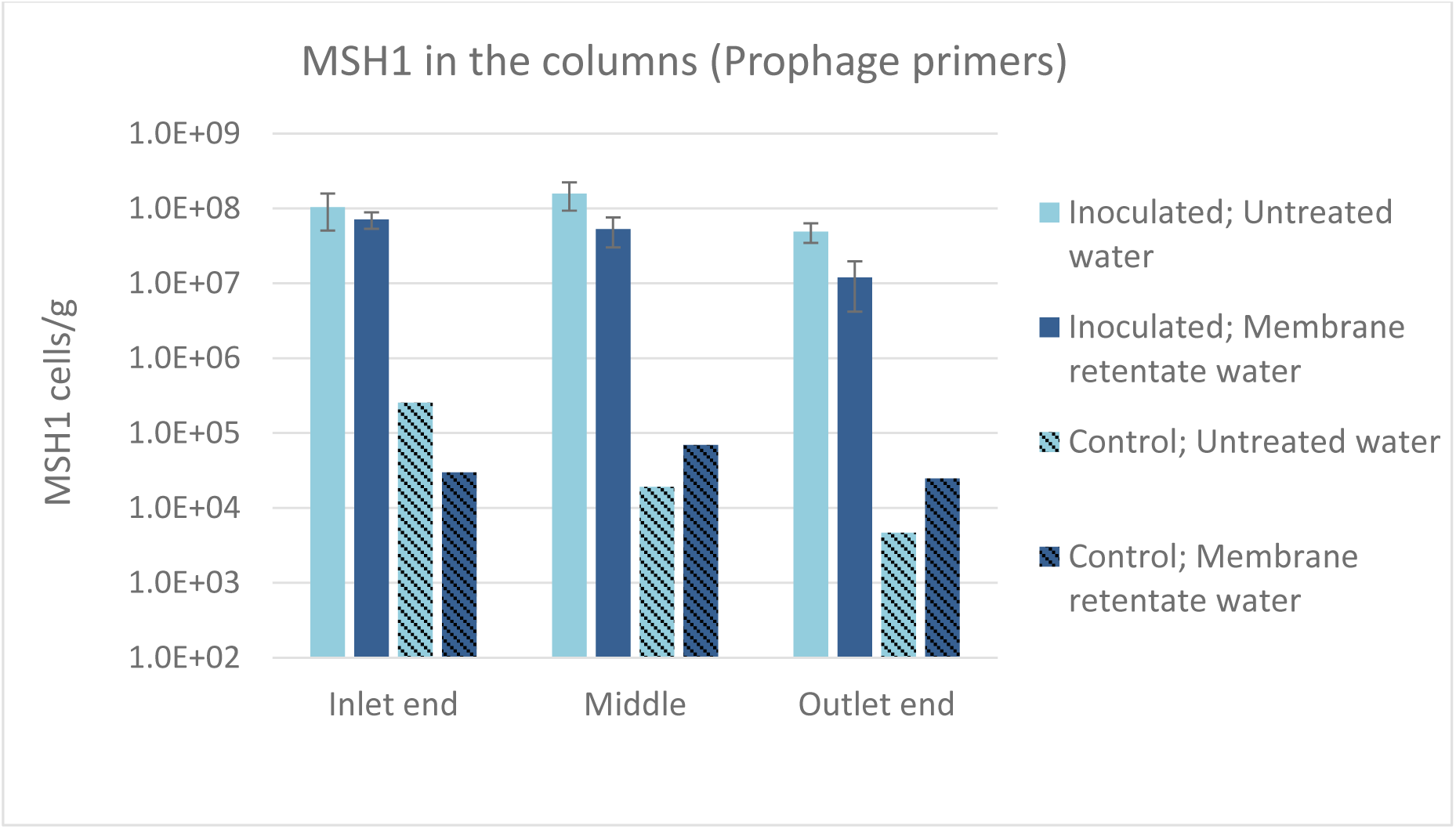
Number of *Aminobacter* sp. MSH1 cells (determined by qPCR applying the newly designed specific primers) detected in the sand sampled from different locations within the columns at experimental termination. Columns received during operation a feed of either untreated water with 0.3 µg l^-1^ BAM (light blue) or membrane retentate water with 3.0 µg l^-1^ BAM (dark blue bars). For the columns inoculated with *Aminobacter* sp. MSH1 data are presented as means (n = 3) with error bars showing standard error. Two control columns were run without initial inoculation (hatched bars; n = 1).

**Fig. S2:**
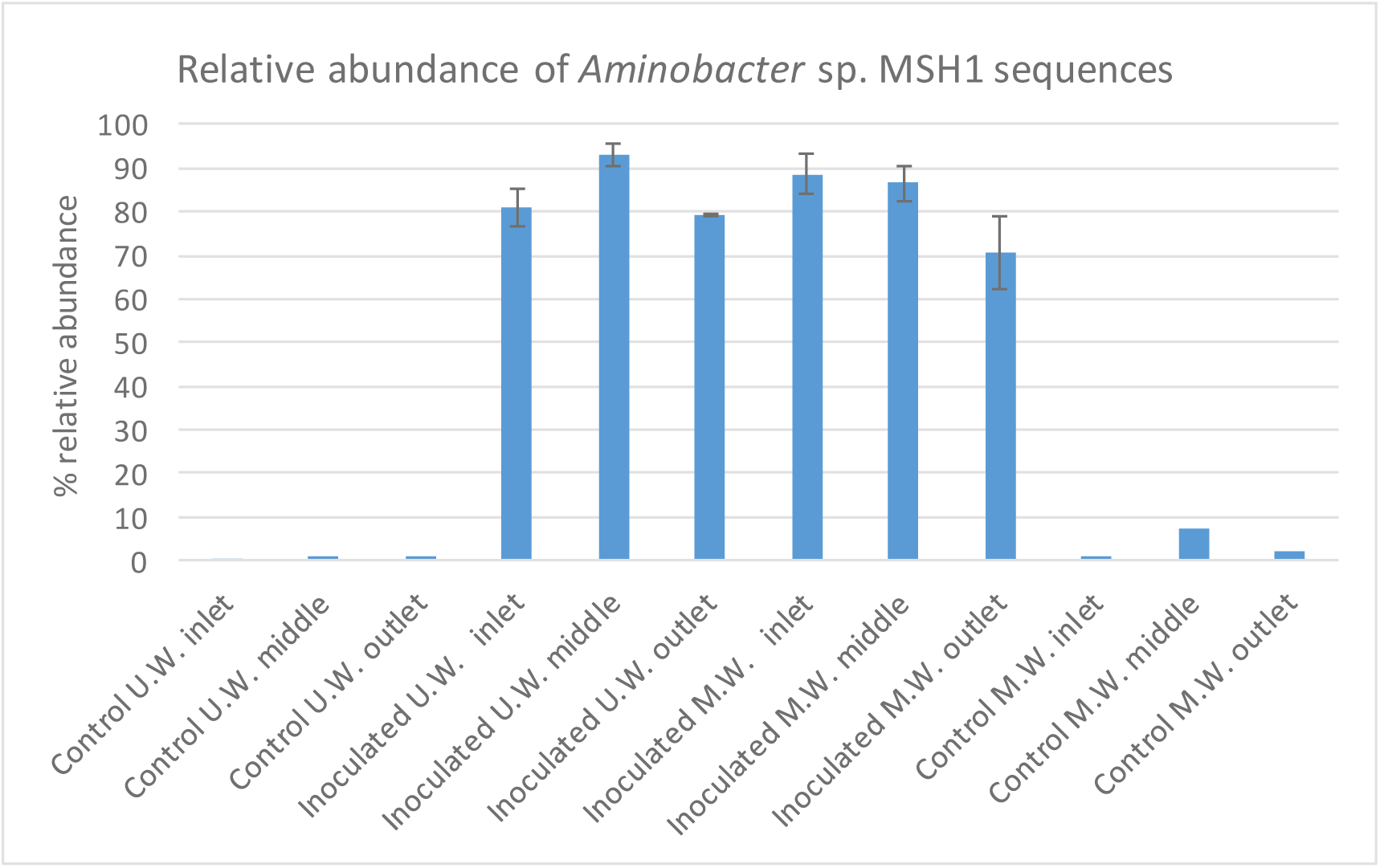
the relative abundance of sequences belonging to *Aminobacter* sp. MSH1 (i.e. Sequences with ≥ 97% similarity to *Aminobacter* sp. MSH1) in the complete microbial communities of the columns.

**Fig. S3.**
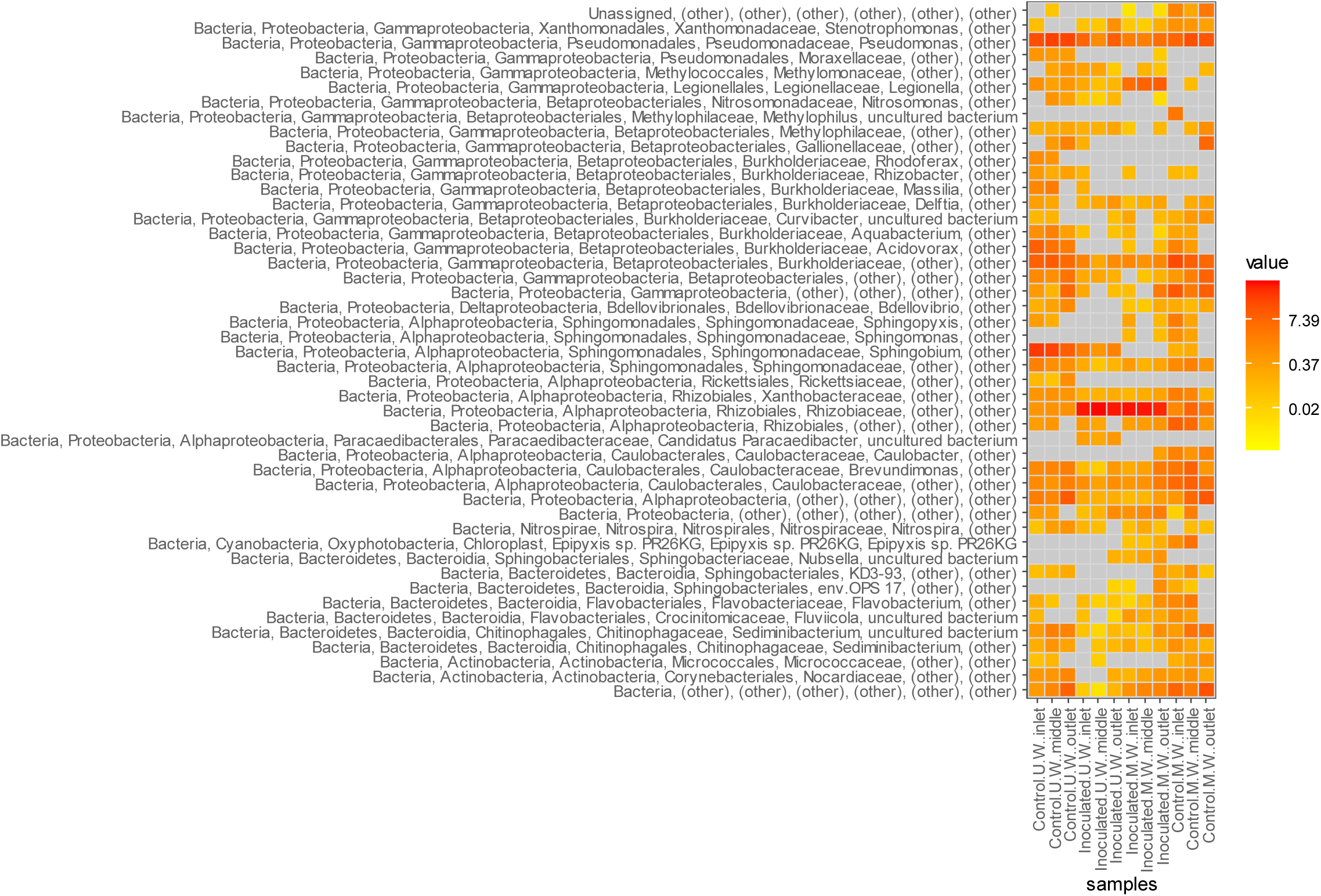
Relative abundance of the OTUs found in the prokaryotic communities, where sequences with ≥ 97% similarity to *Aminobacter* sp. MSH1 had been removed from the dataset. Only OTUs having an accumulated relative abundance of >1% across all samples are included.

